# Differential effects of commensal bacteria on progenitor cell adhesion, division symmetry and tumorigenesis in the *Drosophila* intestine

**DOI:** 10.1101/799981

**Authors:** Meghan Ferguson, Kristina Petkau, Minjeong Shin, Anthony Galenza, David Fast, Edan Foley

## Abstract

Microbial factors influence homeostatic and oncogenic growth in the intestinal epithelium. However, we know little about immediate effects of commensal bacteria on stem cell division programs. In this study, we examined effects of commensal *Lactobacillus* species on homeostatic, and tumorigenic stem cell growth in the *Drosophila* intestine. We identified *Lactobacillus brevis* as a potent stimulator of stem cell growth. In a wildtype midgut, *Lactobacillus brevis* activates growth regulatory pathways that drive stem cell divisions. In a Notch-deficient background, *Lactobacillus brevis*-mediated growth causes rapid expansion of mutant progenitors, leading to accumulation of large, multi-layered tumors throughout the midgut. Mechanistically, we showed that *Lactobacillus brevis* disrupts expression and subcellular distribution of progenitor cell integrins, supporting symmetric divisions that expand intestinal stem cell populations. Collectively, our data emphasize the impact of commensal microbes on growth and maintenance of the intestinal progenitor compartment.

## INTRODUCTION

Multipotent intestinal stem cells (ISCs) divide at a rate that matches the loss of dead or damaged epithelial cells, maintaining an effective barrier to invasion by gut-resident microbes (Alam and Neish, 2018; Amcheslavsky et al., 2009; Biteau and Jasper, 2011; Buchon et al., 2009a; Jiang et al., 2009; Shaw et al., 2010). Commensal bacteria promote epithelial growth (Broderick et al., 2014; Buchon et al., 2009b; Cheesman et al., 2011; Li et al., 2012; Zackular et al., 2013), and disruptions to microbiota composition or diversity are associated with proliferative diseases such as colorectal cancer (Gagnière et al., 2016; Zeller et al., 2014). Thus, we consider it important to understand how the gut bacterial community influences ISC growth and differentiation.

*Drosophila melanogaster* is a popular system to study microbial control of ISC growth due to an extensive toolkit for host genetic manipulation, and a simple, cultivable microbiome that is easy to modify (Broderick and Lemaitre, 2012; Koyle et al., 2016). Importantly, key regulators of ISC growth are evolutionarily conserved between flies and vertebrates (Miguel-Aliaga et al., 2018). For example, vertebrate and fly ISCs reside in a niche that uses related growth factors to direct stem cell division and differentiation (Jiang and Edgar, 2011; Morrison and Spradling, 2008; Takashima and Hartenstein, 2012). Integrins are particularly important niche regulators of ISC division. Integrins anchor fly ISCs to a basal extracellular matrix, orienting the mitotic spindle at an angle to the basement membrane, and ensuring polarized distribution of cell fate determinants (Chen et al., 2018; Goulas et al., 2012). In asymmetric divisions, the apical daughter exits the niche and terminally differentiates as a mature epithelial cell (Martin et al., 2018; Micchelli and Perrimon, 2006; Ohlstein and Spradling, 2006; Ohlstein and Spradling, 2007), while the basal daughter remains within the niche, where it retains ‘stemness’. Depletion of integrins from the fly midgut diminishes asymmetric division frequency, promoting symmetric expansion of stem cell lineages and epithelial dysplasia (Goulas et al., 2012; Lin et al., 2013; Okumura et al., 2014). Notably, relationships between integrins and ISC growth are evolutionarily conserved, as integrin loss also causes hyperplasia in the mouse intestine (Jones et al., 2006).

In flies, most asymmetric divisions generate a post-mitotic enteroblast that differentiates as a large, absorptive enterocyte in response to Notch pathway signals (Biteau and Jasper, 2014; Guo and Ohlstein, 2015; Ohlstein and Spradling, 2007; Zeng and Hou, 2015). Loss of Notch from stem cell/enteroblast progenitor pairs leads to rapid growth of epithelial tumors characterized by hyperplastic stem cells, absence of enterocytes, and accumulation of secretory enteroendocrine cells (Patel et al., 2015). Disruptions to Notch cause similar dysplastic phenotypes in fish and mice (Crosnier et al., 2005; Qiao and Wong, 2009), and spontaneous accumulation of mutations at the *Notch* locus is linked to age-dependent development of intestinal tumors in adult *Drosophila* (Siudeja et al., 2015).

Previous work established that pathogenic *Pseudomonas aeruginosa* causes tumors in flies that express a latent Ras oncogene in progenitor cells (Apidianakis et al., 2009). More recently, we used the Notch-deficient model to show that commensal bacterial are also effective agents of intestinal tumorigenesis (Petkau et al., 2017). However, it is unclear which taxa promote tumors, and how this happens. To understand how gut bacteria regulate epithelial growth, we determined the impacts of common fly commensals on wildtype and Notch-deficient progenitors. We found that *Lactobacillus brevis* stimulated stem cell growth, regardless of host genotype, while a close relative, *Lactobacillus plantarum*, did not. Upon further analysis, we found that the *L. brevis* cell wall was sufficient to promote tumors in Notch-deficient intestines. Mechanistically, *L. brevis* decreased expression, and altered the subcellular distribution of progenitor cell integrins. Consistent with critical requirements for integrins in regulating asymmetric progenitor growth, we found that association with *L. brevis* increased the frequency of symmetric stem cell divisions, resulting in greater numbers of ISCs than post-mitotic enteroblasts. Combined, our data implicate a common fly commensal in the symmetric expansion of stem cell lineages, promoting growth of stem cells that harbor tumorigenic lesions.

## RESULTS

### *L. brevis* promotes tumor growth in the *Drosophila* intestine

To study bacterial effects on intestinal tumors, we used the temperature-controlled *escargot-GAL4, GAL80^ts^, UAS-GFP* (*esg^ts^*) transgenic fly line to express an inducible *Notch* RNAi construct (*UAS-N^RNAi^*) in ISCs and enteroblasts (collectively referred to as progenitor cells) at the restrictive temperature of 29°C. Intestines of control *esg^ts^/+* females contained evenly distributed GFP-positive progenitors and prospero-positive enteroendocrine cells in a simple epithelium dominated by large, polyploid enterocytes (Fig. 1B, Fig. S1A). In line with an earlier study that described massive stem cell growth in flies with *Notch*-deficient progenitors (Patel et al., 2015), we found that depletion of *Notch* (*esg^ts^/N^RNAi^*) caused multilayered midgut tumors populated by excess progenitor and enteroendocrine cells within five days (Fig. 1C, Fig. S1B). As tumors were evident in female intestines, but largely absent from males (Fig. S1C), we performed all subsequent experiments on adult female posterior midguts.

**Figure 1.**
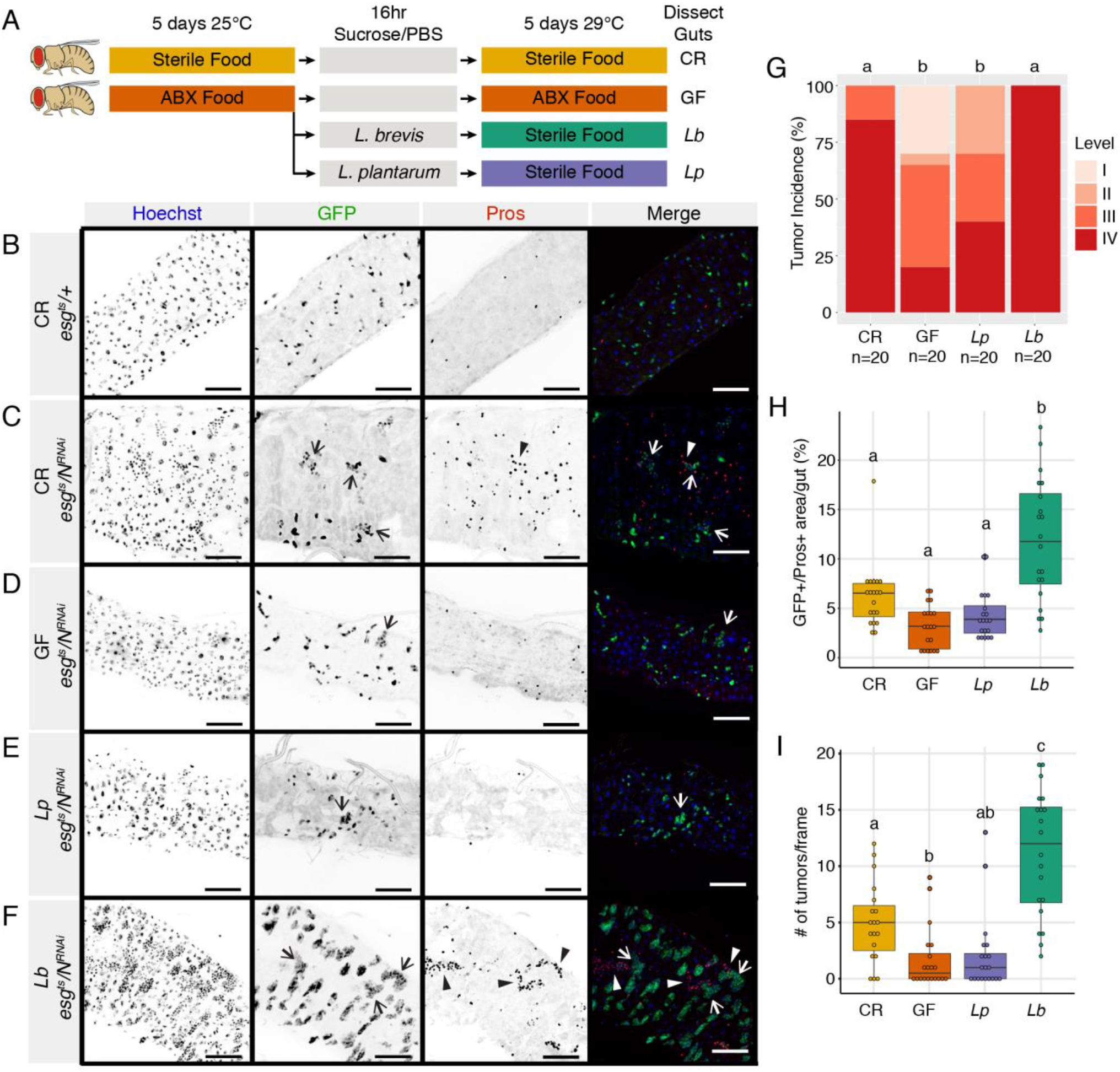
*L. brevis* promotes tumor growth in the *Drosophila* intestine. **(A)** Scheme for generating GF and gnotobiotic flies alongside CR controls. ABX = food with antibiotic cocktail **(B)** Images of wild-type CR *esg^ts^/+* intestines **(C-F)** Images of *esg^ts^/N^RNAi^* posterior midguts 5 days after *Notch* knockdown and microbial manipulation. Hoechst marks DNA (blue), GFP marks esg+ progenitor cells (green) and Pros marks enteroendocrine cells (red). Level III tumours (open arrowheads) and level IV tumours (closed arrowheads). Scale bars = 50μm. **(G)** Tumor incidence in CR, GF, *Lp* and *Lb* monoassociated intestines after 5 days of *Notch* depletion. Different letters denote significant difference of level IV tumor incidence of p<0.01 with Chi-squared test **(H)** Tumor burden in CR, GF, *Lp* and *Lb* monoassociated intestines after 5 days of Notch depletion. Burden is calculated as the percent area of the intestine that is GFP+ and Pros+. **(I)** Number of tumors per frame of the posterior midgut after 5 days of microbial manipulation and Notch depletion. Different letters in H-I denote significant difference of p<0.05 with ANOVA followed by multiple pairwise Tukey tests.

In contrast to intestines with a conventional microbiome, Notch-deficient tumors rarely appeared in age-matched, germ-free (GF) flies, indicating microbial requirements for tumor growth (Fig. 1D), although we cannot exclude the possibility that tumors eventually form in GF flies with age. To identify bacterial species that promote tumors, we examined posterior midguts of adult *esg^ts^/N^RNAi^* flies that we associated exclusively with common species of *Lactobacillus* commensals, a dominant genus within the fly microbiome (Adair et al., 2018; Wong et al., 2013). To focus exclusively on adult tumors, we raised *esg^ts^/N^RNAi^* larvae with a conventional microbiome under conditions that prevent Notch inactivation. Upon eclosion, we fed adults an antibiotic cocktail that depleted the bacterial microbiome below detectable levels, and re-associated flies with *Lactobacillus brevis* (*Lb*), or *Lactobacillus plantarum* (*Lp*) (Fig. 1A). We compared each mono-association to conventionally reared (CR) *esg^ts^/N^RNAi^* flies that contained a poly-microbial gut microbiota. Mono-association of *esg^ts^/N^RNAi^* flies with *Lp* resulted in few visible tumors (Fig. 1E). In contrast, mono-association with *Lb* caused multiple, large tumors throughout the posterior midgut (Fig. 1F), indicating that *Lb* is sufficient for tumor development.

We then quantified impacts of bacterial association on midgut tumors. First, we developed a four-point system to classify intestines, ranging from no visible defects (level I) to intestines with progenitor and enteroendocrine-rich tumors (level IV, Fig. S1D). In a blinded assay, we categorized 85% of CR *esg^ts^/N^RNAi^* intestines as level IV, whereas only 20% of GF intestines belonged to the same category (Fig. 1G), confirming bacterial effects on gut tumors. Consistent with our initial observations, GF and *Lp*-associated intestines had similarly mild levels of midgut tumors (Fig. 1G). In contrast, all intestines associated with *Lb* had level IV tumors within five days of Notch inactivation. To measure total tumor size per midgut, we quantified the posterior midgut area occupied by progenitor and enteroendocrine cells in the respective groups. Association with *Lb* significantly enhanced accumulation of progenitors and enteroendocrine cells in *esg^ts^/N^RNAi^* intestines compared to CR, GF or *Lp* mono-associated flies, supporting a role for *Lb* in promoting tumors (Fig. 1H). To determine if the larger tumor areas in *Lb-*associated flies are a result of increased tumor initiation, or accelerated tumor growth we quantified numbers of tumors in each intestine. Similar to our assessment of tumor size, association with *Lb* had a significant impact on tumor numbers, resulting in approximately three times as many tumors per gut as CR counterparts (Fig. 1I). Collectively, our data indicate that association with *Lb* increases the frequency of midgut tumor initiation.

To determine which factors from *Lb* promote tumors, we measured tumors in flies that we continuously fed heat-killed *Lb* or cell wall derived from *Lb* for five days. GF intestines fed heat-killed *Lb* had similar tumor levels (Figure 2A), and similar progenitor and enteroendocrine cell expansions as flies fed live *Lb* (Fig. 2B), indicating that bacterial metabolites are not required for tumors. Instead, GF *esg^ts^/N^RNAi^* flies fed *Lb* cell wall extract had similar tumor levels (Fig. 2C), and similar increases in progenitor and enteroendocrine cell numbers as GF counterparts fed dead *Lb* (Fig. 2D). Finally, we noticed that *esg^ts^/N^RNAi^* flies mono-associated with *Lb* died significantly faster than CR counterparts, while *Lb* did not shorten the lifespan of *esg^ts^/+* controls (Fig. S2), arguing that cell wall components from *L. brevis* promote initiation of Notch-deficient tumors, resulting in premature host death.

**Figure 2.**
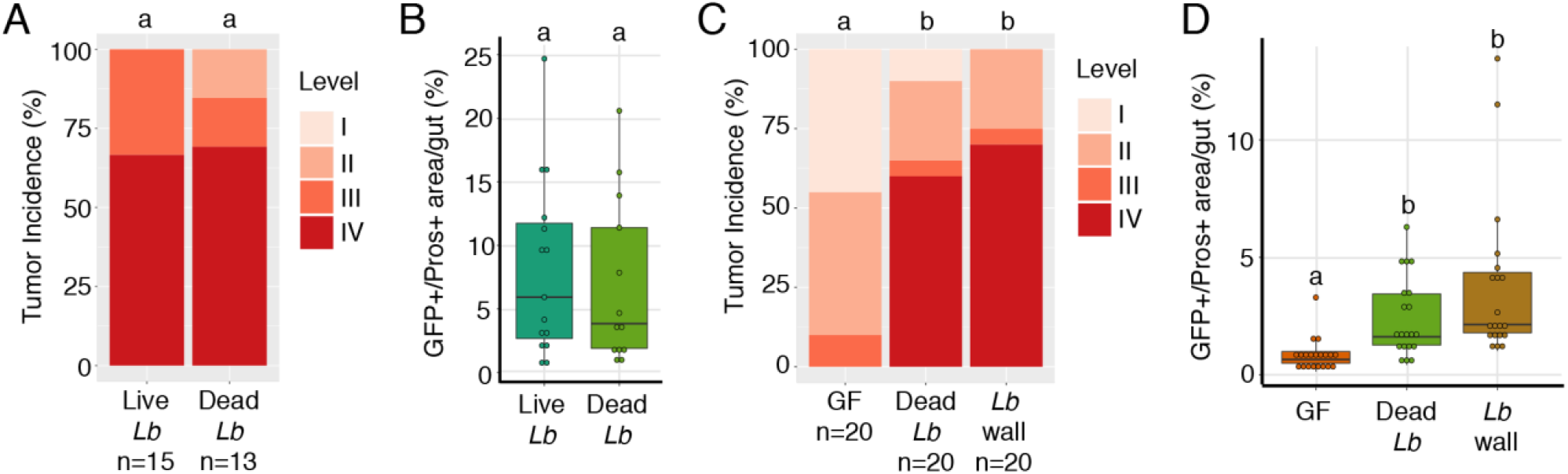
*L. brevis* cell wall is sufficient to promote tumor growth. Tumor incidence **(A)** and burden **(B)** of *esg^ts^/N^RNAi^* flies colonized with live *Lb* or GF *esg^ts^/N^RNAi^* fed heat killed *Lb* mixed into sterile food. Same letters denote no significant difference between level IV tumor incidence at p = 0.05 with Chi squared test **(A)** or tumor burden using pairwise Wilcoxon test **(B).** Tumor incidence **(C)** and burden **(D)** of GF *esg^ts^/N^RNAi^* flies fed heat killed *Lb* or cell wall extract in PBS/sucrose on filter paper. Different letters denote significant difference of p < 0.05 with Chi squared test **(C)** or pairwise Wilcoxon test **(D).**

### Notch inactivation modifies expression of growth, differentiation, and immunity regulators in progenitors

To determine how *Lb* affects Notch-deficient progenitors, we used RNA-sequencing to identify the transcriptional profiles of FACS-purified, GFP-positive progenitors from *Lb*-associated *esg^ts^/+* and *esg^ts^/N^RNAi^* intestines (Fig. 3A). As controls, we sequenced transcriptomes of *esg^ts^/+* and *esg^ts^/N^RNAi^* progenitors from GF flies, or flies that we mono-associated with *Lp*. Principal Component Analysis (PCA) revealed that Notch-deficient progenitors segregate from wildtype progenitors along PC1, regardless of bacterial association (Fig. 3B). Differential gene expression analysis showed that the majority of gene expression profiles altered by Notch-depletion were shared between GF, *Lb*-associated and *Lp*-associated intestines (Fig. 3C), indicating the existence of a microbe-independent core response to loss of Notch in progenitors. GO term analysis of the core Notch-deficient response revealed significant upregulation of biological processes involved in cell division (Fig. 3D), and diminished expression of Notch-responsive *Enhancer of split (E(spl))* complex genes required for enteroblast differentiation (Fig. 3E). In addition to effects on growth and differentiation, we observed unexpected downregulation of immune pathway regulators in progenitors that lacked *Notch* (Fig. 3D-E). Decreased expression of immune regulators is not secondary to tumor development, as we saw similar changes in intestines of GF, and *Lp*-associated flies. In each case, Notch inactivation diminished expression of essential components of the antibacterial Immune Deficiency (IMD) pathway such as *imd*, *IKKβ*, and *Rel*, as well as prominent IMD response genes such as *pirk*, and multiple PGRP family members (Fig. 3E). These data suggest a genetic link between Notch signaling and antibacterial responses in the progenitor compartment, and match previous reports that mutation or activation of the IMD pathway alters expression of Notch pathway genes in the fly intestine (Broderick et al., 2014; Petkau et al., 2017).

**Figure 3.**
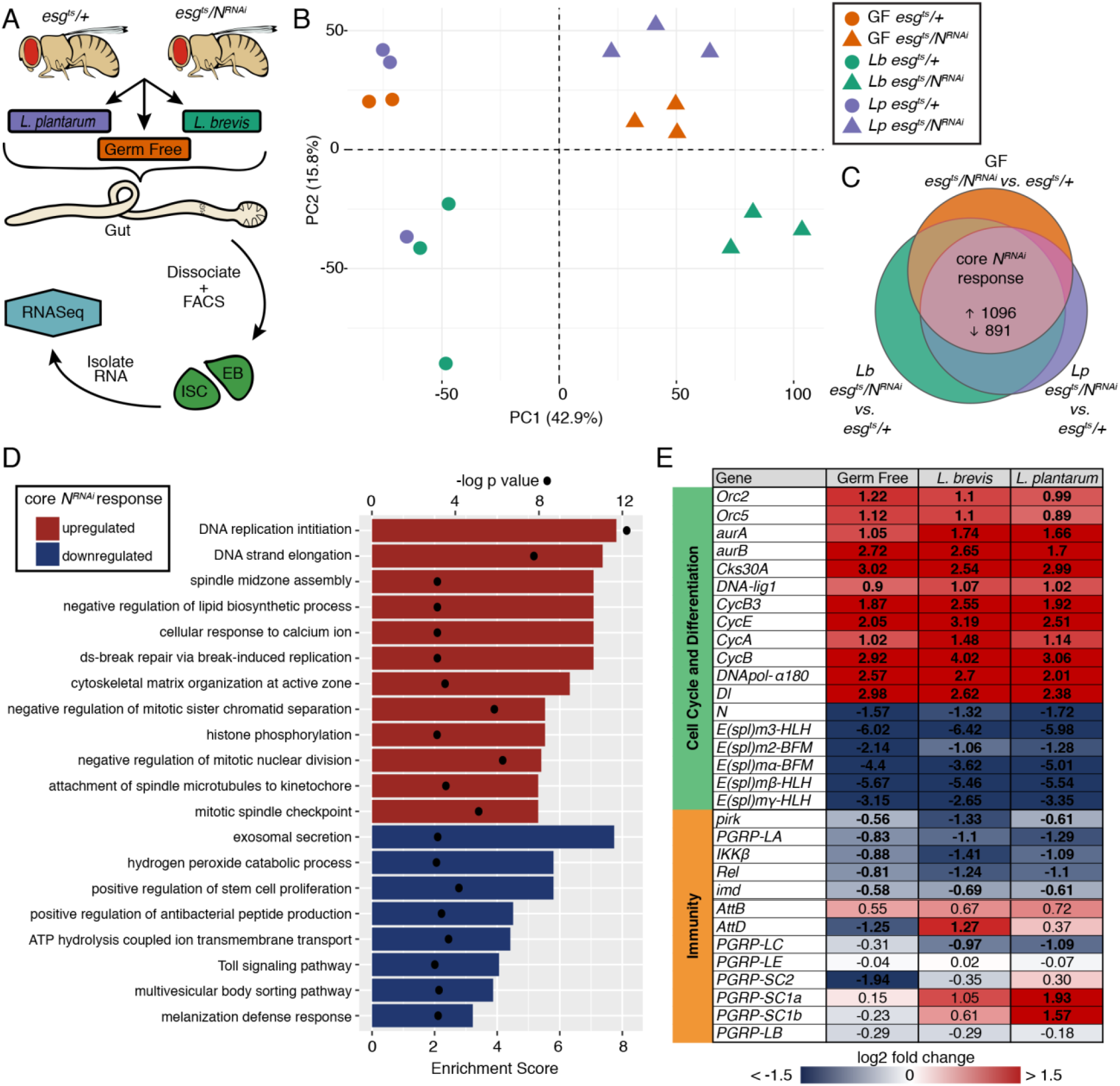
Notch inactivation decreases expression of immunity regulators in intestinal progenitors. **(A)** Workflow for the RNA-Seq of intestinal progenitors upon *Notch* knockdown and *Lb* colonization. **(B)** PCA plot from RNA-Seq project. Circles represent *esg^ts^/+* and triangles represent *esg^ts^/N^RNAi^* replicates. Different colors represent GF (orange), *Lb* (teal) or *Lp* (purple). **(C)** Genes altered by *Notch* knockdown (p<0.01, FDR <5%) in each microbial context showing the core response to knockdown of *Notch*. **(D)** Biological process GO terms enriched in the core *Notch* response. Enrichment score shown as bars and p values shown as dots. **(E)** Log2 fold change of *Notch* response genes involved in cell cycle/differentiation and immunity. Values in bold are significantly altered genes with a p<0.01 and FDR 5%. All genes above the double line are part of the core Notch response whereas genes below (*AttB* to *PGRP-LB*) are immune genes not included in the core response. Each column is a direct comparison of *esg^ts^/N^RNAi^* to *esg^ts^/+* under GF, *Lb* or *Lp* conditions.

### Notch-deficiency promotes intestinal association with *L. brevis*

As tumor growth frequently involves shifts in microbiota composition, we determined effects of Notch inactivation on host association with *Lb* and *Lp*. First, we measured the intestinal bacterial load of *esg^ts^/+* and *esg^ts^/N^RNAi^* flies that we mono-associated with the respective strains for five days. In these experiments, we quantified bacterial load in the same cohort of flies that we used to measure tumors in Figure 1, allowing us to determine if host-microbe associations correlate with midgut tumors. For *Lb* and *Lp*, we observed significantly increased bacterial loads in Notch-deficient intestines compared to wild-type controls (Figure 4A), suggesting effects of host genotype on bacterial association. Importantly, there were no differences between *Lp* or *Lb* loads in Notch-deficient guts. Thus, the identity of associated bacteria, not the abundance, determines tumors in the host.

**Figure 4.**
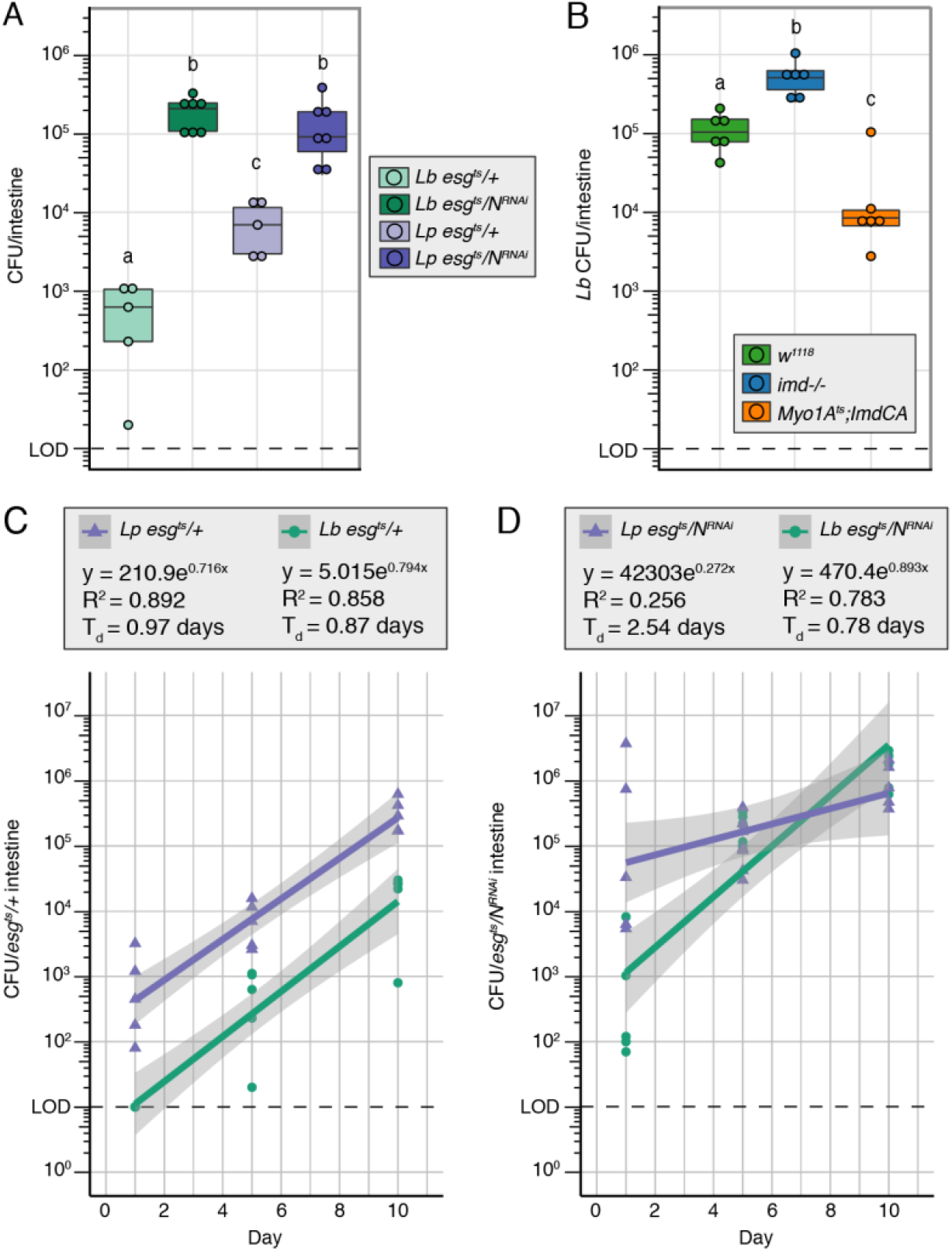
Notch-deficiency promotes intestinal association with *L. brevis*. **(A)** Colony forming units (CFU) of *Lb* (green) and *Lp* (purple) per fly intestine in monoassociated *esg^ts^/+* and *esg^ts^/N^RNAi^* 5 days after transgene expression and bacterial colonization. **(B)** CFU of *Lb* per fly intestine 10 days after transgene expression/bacterial colonization in *Lb* monoassociated wild-type (*w^1118^*), *imd-/-*, and *Myo1A^ts^;ImdCA*. For **(A)** and **(B),** different letters denote significance at p<0.05 with multiple pairwise Wilcoxon tests. **(C)** CFU of *Lb* (green) and *Lp* (purple) over time in *esg^ts^/+* intestines. **(D)** CFU of *Lb* (green) and *Lp* (purple) over time in *esg^ts^/N^RNAi^* intestines. For **(C)** and **(D)**, x axis is days post transgene expression/bacterial colonization and line represents exponential trendline with shaded region being a 95% confidence interval. T_d_ = doubling time. LOD = Limit of detection.

As bacterial association is higher in Notch-deficient intestines than wildtype intestines (Fig. 4A), and Notch inactivation diminishes expression of IMD pathway components, we asked if IMD affects host association with *Lb*. Consistent with a role for IMD in the control of intestinal *Lb*, we found that *imd* mutants had significantly higher *Lb* loads than wild-type controls ten days after mono-association with *Lb* (Fig. 4B). Conversely, constitutive activation of IMD in enterocytes (*Myo1A^ts^*/*ImdCA*) reduced *Lb* load to approximately 4% of that found in *imd* mutants (Figure 4B). These data support a role for IMD in regulation of intestinal *Lb*. However, it is important to note that the increased bacterial abundance in *imd* mutants is considerably less pronounced than increases observed upon *Notch* depletion (compare Fig. 4A and 4B). Thus, we believe that additional, IMD-independent mechanisms control bacterial numbers in *esg^ts^/N^RNAi^* intestines that require identification.

Finally, we measured the effects of Notch inactivation on host association with *Lactobacillus* commensals. Here, we completed a longitudinal measurement of bacterial load in intestines of *esg^ts^/+* and *esg^ts^/N^RNAi^* flies that we mono-associated with *Lp* or *Lb*. In general, our data match earlier reports that total numbers of intestinal bacteria increase in flies with age (Clark et al., 2015; Guo et al., 2014). In wild-type *esg^ts^/+* intestines, the rates of increase in host-association with *Lp* and *Lb* are nearly indistinguishable, with *Lb* associating to lower levels at all times tested (Fig. 4C). Initially, *Lb* also associated with *esg^ts^/N^RNAi^* intestines to lower levels than *Lp*. However, we noted substantial effects of Notch inactivation on subsequent progressions in host-microbe association. In this case, association with *Lp* increased at a considerably slower rate than association with *Lb* (Fig. 4D). Exponential regression analysis revealed that host-associated *Lb* loads double at similar rates in intestines of *esg^ts^/+* (0.87d) and *esg^ts^/N^RNAi^* (0.78d) flies. In contrast, host-associated-*Lp* loads double at a considerably slower rate in N-deficient intestines (2.54d, Fig. 4D), than wild-type intestines (0.97d, Fig. 4C), suggesting that Notch knockdown hinders host association with *Lp*, but has minimal effects on association with *Lb*.

### *L. brevis* decreases expression of integrins in progenitor cells

As *Lb* grows effectively in Notch-deficient intestines, where it promotes tumors, we reasoned that *Lb* will have distinct growth-enhancing effects on progenitor cells. To test this hypothesis, we looked for progenitor cell transcriptional events that were specific to association with *Lb*. Principle component analysis identified a transcriptional response that is unique to *Lb* in wild-type and *Notch*-deficient progenitors (Fig. 2B, Fig. S3A,B). Regardless of host genotype, association with *Lb* specifically increased expression of genes required for cell growth, such as DNA replication, and mitotic spindle organization (Fig. 5A), as well as prominent cell cycle and growth regulators (Fig. 5B). We noticed a particularly striking inhibitory effect of *Lb* on expression of genes involved in cell-cell adhesion, cell-matrix adhesion and cell polarity (Fig. 5A), especially genes that encode integrin complex proteins (Fig. S3C). For example, association with *Lb* led to diminished expression of the alpha and beta-integrins *scab* and *myospheroid* (*mys*), the talin ortholog, *rhea*, and the integrin extracellular matrix ligand, *LanA* (Fig. 5B). The effects of *Lb* on expression of genes associated with stem cell adhesion were independent of host genotype, as we observed the same phenotypes in progenitors of *Lb-*associated *esg^ts^/+* and *esg^ts^/N^RNAi^* flies (Fig. 5B).

**Figure 5.**
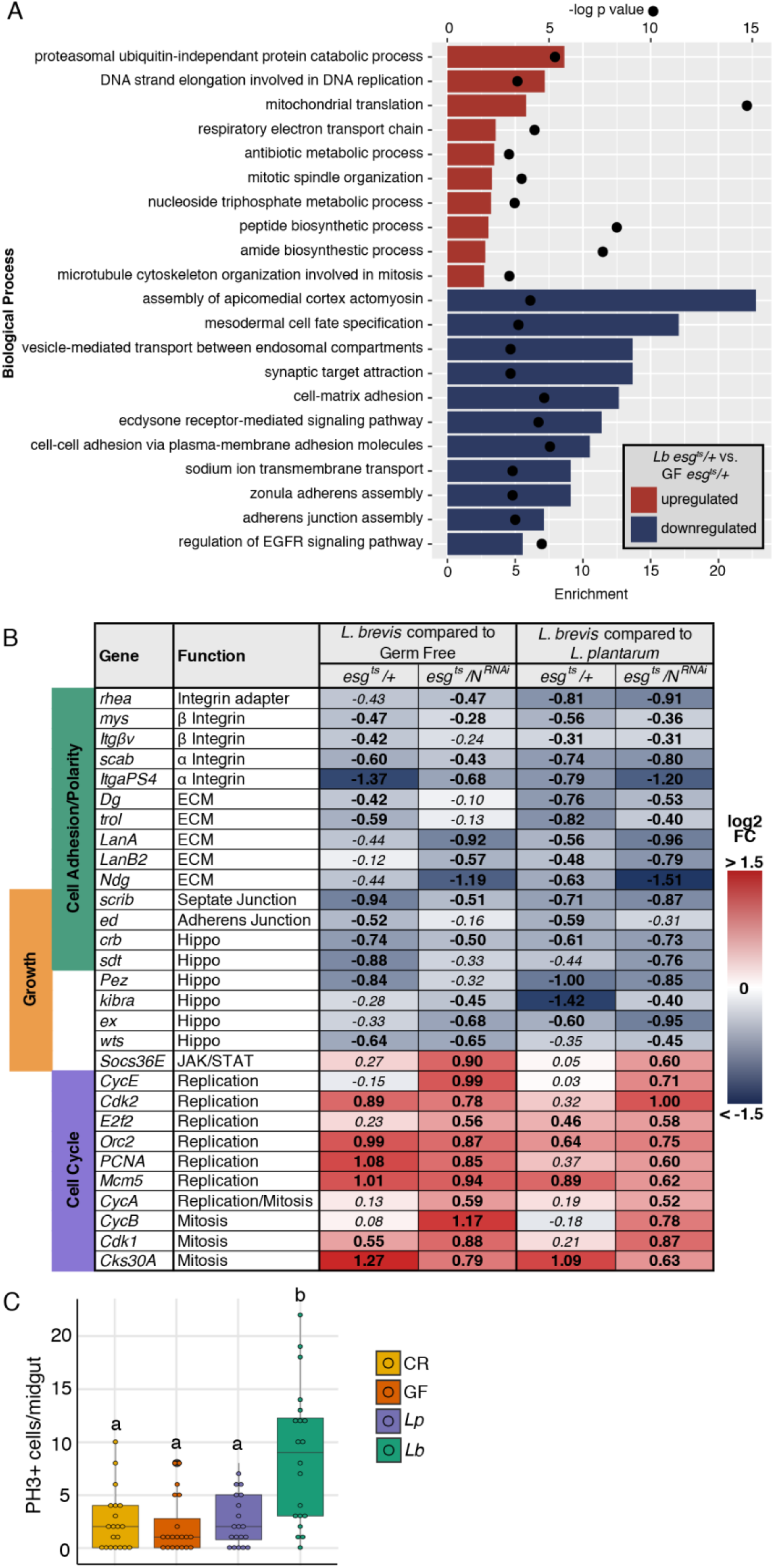
*L. brevis* decreases expression of integrins in progenitor cells. **(A)** Biological process GO terms enriched in progenitors from *esg^ts^/+* colonized with *Lb* compared to GF. Enrichment score shown as bars and p values shown as dots. **(B)** Log2 fold change of genes involved in cell adhesion/polarity, Growth and Cell cycle affected by *Lb* in comparison to either GF or *Lp* colonization in *esg^ts^/+* and *esg^ts^/N^RNAi^* progenitors. Bolded values are those with a p value <0.05 and FDR<5%. **(C)** Number of PH3+ cells per *esg^ts^/+* midgut in gnotobiotic flies 8 days after colonization. Different letters denote significant difference of p<0.01 by multiple pairwise Wilcoxon tests.

Given the positive effects of *Lb* on expression of growth regulators in *esg^ts^/+* and *esg^ts^/N^RNAi^* progenitors, we asked if *Lb* activates ISC division in wild-type progenitors. To answer this question, we mono-associated GF wild-type (*esg^ts^/+*) flies with *Lb* and quantified phospho-histone 3-positive (PH3+) mitotic cells in adult midguts. Similar to effects on tumors, *Lb* stimulated growth of wildtype progenitors to significantly higher levels than CR, GF, or *Lp*-mono-associated flies (Fig. 5C). Thus, our data indicate that association with *Lb* diminishes expression of genes required for progenitor adhesion to the extracellular matrix, and induces expression of genes required for epithelial growth, promoting ISC division in wildtype and Notch-deficient progenitors.

### *L. brevis* colonization disrupts integrin localization independent of division

We were particularly intrigued by effects of *Lb* on expression of integrins that anchor progenitors within the niche. Therefore, we asked what effects *Lb* has on progenitor cell adhesion and morphology. In a preliminary experiment, we used transmission electron microscopy to visualize posterior midguts of CR, and *Lb-*associated wild-type flies. CR intestines contained basal progenitors in close association with the extracellular matrix (Fig. 6A, B, Fig. S4A, B for additional examples). Mono-association with *Lb* appeared to disrupt intestinal organization, generating round progenitors that shifted apically relative to the extracellular matrix, and lacked discernible contact with larger enterocytes or extracellular matrix (Fig. 6C, D, Fig. S4C, D for additional examples). These morphological changes appear specific to progenitors, as no defects were apparent in the shape, or relative position, of surrounding enterocytes. The apparent shift in progenitor localization in *Lb*-associated intestines prompted us to ask if *Lb* modifies integrin distribution. To answer this question, we determined the subcellular localization of the β-integrin, mys, in sagittal sections of GF intestines, or intestines that we associated with *Lb*. In GF *esg^ts^/+* flies, we detected basolateral enrichment of mys in GFP-positive progenitors (Fig. 6E). In contrast, and similar to our electron microscopy results, we found that *Lb* colonization caused progenitors to round up and adopt a more apical position within the epithelium (Fig. 6F). Furthermore, association with *Lb* had visible impacts on mys localization, characterized by discontinuous basolateral distribution, and atypical apical enrichment of mys (Fig. 6F, arrowheads). To directly measure effects of *Lb* on subcellular distribution of integrins, we developed an immunofluorescence-based assay that allowed us to quantify apical:basolateral ratios of mys in progenitors (Fig. S5). With this assay, we detected basal enrichment of mys in GF progenitors (Fig. 6H). Association with *Lb* shifted the distribution of mys, resulting in significant increases in apical mys (Fig. 6H). To determine if *Lb*-dependent effects on integrin subcellular distribution are downstream consequences of stem cell division, we blocked growth in progenitors of *Lb*-associated flies by expressing the cell cycle inhibitor *dacapo* (*esg^ts^/dap*) (Fig. 6I). Notably, when we examined growth-impaired midguts, we found that *Lb* continued to cause increases in apical mys (Fig. 6G-H), indicting that *Lb* alters apicobasal integrin distribution independent of ISC divisions.

**Figure 6.**
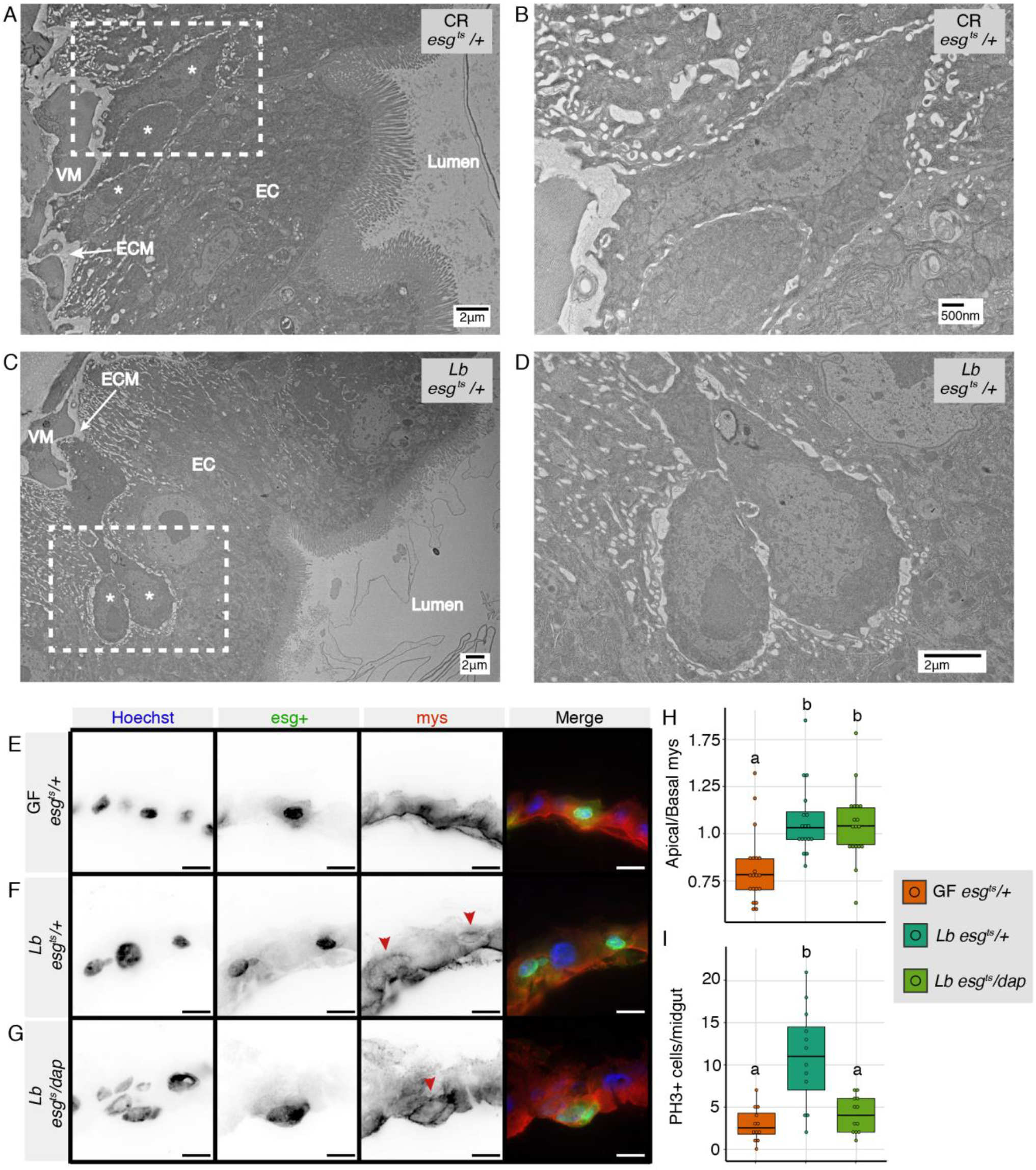
*L. brevis* colonization disrupts integrin localization independent of division. **(A-D)** TEM images of posterior midgut cross sections from *esg^ts^/+* CR (A-B) and *Lb* monoassociated (C-D) intestines. VM = visceral muscle, ECM = Extracellular matrix, EC = Enterocyte, * = progenitors. Dashed boxes in (A) and (C) showed zoomed areas of (B) and (D) respectively. **(E-G)** Immunofluorescence images of posterior midgut sagittal sections from GF (E) and *Lb* (F) monoassociated *esg^ts^/+* and *Lb* monoassociated *esg^ts^/dap* (G) flies after 8 days of transgene expression and bacterial colonization. Hoescht marks DNA (blue), esg marks progenitors (green) and mys marks integrins (red). Top of image is the apical/luminal side, bottom of image is the basal side of the epithelium. Red arrowheads = apical integrin mis-localization. Scale bars = 10μm **(H)** Quantification of apical/basal progenitor cell mys intensity ratio from images captured from conditions in E-G. **(I)** Number of PH3+ cells per midgut of GF and *Lb* monoassociated *esg^ts^/+* and *Lb* monoassociated *esg^ts^/dap* intestines after 8 days of transgene expression/bacterial colonization. For H and I, different letters denote significant difference of p<0.01 with ANOVA followed by multiple pairwise Tukey tests.

### *L. brevis* alters progenitor cell identity and promotes symmetric expansion of stem cell lineages

Loss of intestinal integrins results in aberrant stem cell divisions with substantial effects on organization of the progenitor compartment (Goulas et al., 2012). Therefore, we asked what effects *Lb* has on progenitor cell identity. We first stained intestines of GF flies, or flies that we mono-associated with *Lb* for the ISC marker Delta (Fig. 7A-B). Compared to GF intestines, *Lb* association significantly increased the proportion of Delta+ cells within the *esg*+ progenitor pool (Fig. 7C). In addition, association with *Lb* increased expression of genes involved in stem cell identity, maintenance and differentiation, such as *Dl* and *Enhancer of split (E(Spl))* complex genes (Fig. 7D). As *Lb* also prompts stem cell division (Fig. 5C), we asked what effects *Lb* has on enteroblast proportions. Here, we quantified marker expression in midguts of *esgGAL4, UAS-CFP*, *Su(H)-GFP*; *GAL80^ts^* flies that we raised under conventional conditions, germ-free conditions, or mono-associated with *Lb* (Fig. 7E-G). In this line, stem cells that express the progenitor cell marker *esg* are visible as CFP single-positive cells. In contrast, progenitors that express the enteroblast marker *Su(H)*, are visible as CFP/GFP double-positive cells. We found that CR flies had approximately equal numbers of *Su(H)-*positive and *Su(H)-*negative progenitors, suggesting a 1:1 distribution of stem cells and enteroblasts in midguts of conventional flies (Fig. 7H). Removal of the microbiome increased the proportion of Su(H)+ cells, whereas mono-association with *Lb* had the opposite effect (Fig. 7H). Specifically, we measured a significant decrease in the proportion of Su(H)+ cells within the esg+ population of *Lb*-associated flies (Fig. 7H). Combined with quantification of Dl+ stem cells (Fig. 7A-C), our data indicate that relative to CR or GF flies, *Lb* shifts composition of the progenitor cell compartment towards stem cells.

**Figure 7.**
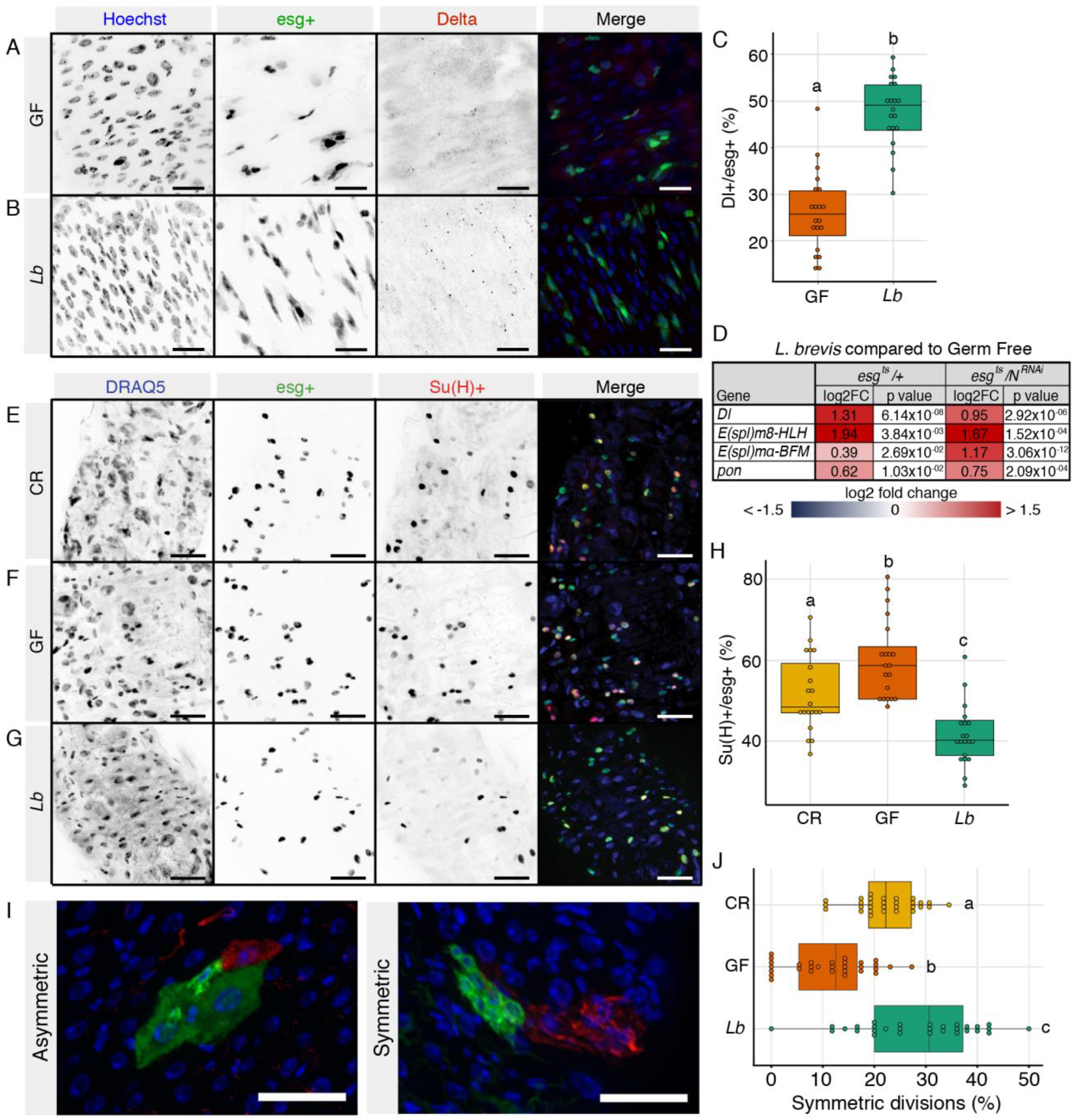
*L. brevis* alters progenitor cell identity and promotes symmetric expansion of stem cell lineages. **(A-B)** Posterior midgut of *esg^ts^/+* GF and *Lb* monoassociated flies where Delta puncta (red) labels presumptive stem cells. Hoescht labels DNA (blue) and esg labels progenitors (green). Scale bars = 15μm **(C)** Percentage of Dl+ cells within the esg+ progenitor population in GF and *Lb esg^ts^/+*. Different letters denote significance of p<0.01 by ANOVA followed by Tukey test. **(D)** Genes differentially expressed in *esg^ts^/+* and *esg^ts^/N^RNAi^* progenitor cells upon *Lb* colonization **(E-G)** Posterior midgut of CR, GF or *Lb* monoassociated *esg^ts^,UAS-CFP,Su(H)-GFP* flies where DRAQ5 labels DNA (blue), esg labels all progenitors (green) and Su(H) labels presumptive enteroblasts (red). Scale bars = 25μm. **(H)** Proportion of Su(H)+ cells within the esg+ progenitor population from CR, GF and *Lb* monoassociated *esg^ts^,UAS-CFP,Su(H)-GFP* intestines. Different letters denote significance at p<0.01 by ANOVA followed by multiple pairwise Tukey tests. **(I**) Representative images of asymmetric and symmetric clones from twin-spot (*hsFLP; FRT40A,UAS-CD2-RFP,UAS-GFP-miRNA/FRT40A,UAS-CD8-GFP,UAS-CD2-miRNA;tubGAL4/+)* flies 8 days after clone induction. Scale bars = 25μm **(J)** Percentage of symmetric divisions in the midgut of CR, GF or *Lb* monoassociated twin-spot flies 8 days after bacterial manipulation and clone induction (n=30). Different letters denote significance at p<0.01 by ANOVA followed by pairwise Tukey tests.

We do not see elevated levels of cell death (Fig. S6A), or increased expression of apoptosis regulators (Fig. S6B) within progenitors of *Lb* mono-associated flies compared to CR or GF controls. Thus, we do not believe that *Lb* affects cell composition within the progenitor compartment by preferentially promoting enteroblast death (Reiff et al., 2019). As an alternative, we tested the hypothesis that *Lb* increases stem cell numbers by promoting symmetric stem cell divisions. Specifically, we used twin-spot Mosaic Analysis with Repressible Cell Markers (MARCM) to visualize and quantify symmetric and asymmetric divisions in intestines of CR, GF, and *Lb* mon-associated flies (Fig. 7I). With these flies, a heat shock induces mitotic recombination and segregates RFP and GFP markers to the two daughter cells (O’Brien et al., 2011; Yu et al., 2009). Asymmetric stem cell divisions generate clones where a single cell of one color resides next to multiple cells of another color. In contrast, symmetric stem cell divisions label approximately equal numbers of intestinal epithelial cells with the respective markers (Fig. 7I). Consistent with earlier reports (Hu and Jasper, 2019; Jin et al., 2017; O’Brien et al., 2011), approximately 23% of all clones were products of symmetric divisions in CR flies (Fig. 7J). Removal of the microbiome resulted in an increase of asymmetric clones (Fig. 7J), indicating that gut bacteria promote symmetric stem cell divisions. Importantly, mono-association with *Lb* significantly increased the frequency of symmetric divisions compared to either GF or CR flies (Fig. 7J). In sum, association with *Lb* resulted in increased symmetric stem cell divisions, expanded stem cell numbers, and decreased enteroblast numbers without progenitor cell death. Combined, our data argue that *Lb* promotes symmetric divisions that expand ISC lineages within the midgut.

## DISCUSSION

Excess microbiota-responsive growth supports development of inflammatory diseases, and hyperplastic expansion of cells that bear oncogenic lesions. To understand how gut bacteria cause progenitor dysplasia, we measured growth in adult *Drosophila* intestines that we mono-associated with common *Lactobacillus* commensals. We focussed on *L. brevis* and *L. plantarum*, as they have established roles in *Drosophila* intestinal homeostasis (Combe et al., 2014; Fast et al., 2018; Iatsenko et al., 2018; Jones et al., 2013; Jones et al., 2015; Lee et al., 2013; Reedy et al., 2019; Storelli et al., 2011). In general, our observations match literature that highlight distinct, context-dependent effects of *Lactobacillus* commensals on juvenile growth, intestinal physiology, and adult longevity. We identified *L. brevis* as a potent stimulator of intestinal tumors. Interestingly, *L. plantarum* did not promote tumor growth even though it grew to similar levels in the intestine as *Lb*, suggesting that *Lp* either fails to stimulate, or actively inhibits, ISC growth. We do not fully understand how the host distinguishes *Lactobacillus* species, although bacterial metabolites are interesting candidates. For example, *Lp* promotes progenitor growth in larvae by stimulating ROS generation in enterocytes (Jones et al., 2013; Jones et al., 2015; Reedy et al., 2019). Likewise, *Lb*-derived uracil activates epithelial generation of ROS in adults, a noxious agent that promotes tissue damage and repair (Lee et al., 2013). In addition, cell walls of *Lp* contain DAP-type peptidoglycan that activates the IMD pathway with substantial effects on transcription (Broderick et al., 2014; Lesperance and Broderick, 2020). In the future, it will be of interest to test the relationships between microbe-specific immune responses, ROS production, and epithelial growth. As poly-bacterial communities have distinct effects on host phenotypes (Gould et al., 2018), it will also be of interest to test effects of interactions among *Lactobacillus* species on epithelial growth and tumorigenesis.

Quantification of host-associated bacteria suggest that physiological disruptions associated with *Notch*-deficiency do not impair accumulation of *Lb* in the gut, which promotes continued growth of *Notch*-deficient tumors. A similar feed-forward loop was described recently in a BMP-deficient tumor model (Zhou and Boutros, 2020), suggesting a conserved relationship between tumor growth and microbiota expansion in flies. Our transcriptional data raise the possibility that inactivation of Notch partially increases *Lb* loads by suppressing expression of IMD pathway regulators. However, given the massive increase in bacterial numbers observed upon Notch knockdown compared to the more moderate increases observed associated with *imd* mutants, we hypothesize that additional mechanisms of Notch-dependant bacterial control exist. As Notch is required for intestinal differentiation, we suggest that a mis-differentiated epithelium is incapable of fully limiting bacterial accumulation in the intestine. For example, Notch inactivation changes the composition, and possibly also gene expression, of differentiated cells within the midgut. Furthermore, Notch controls differentiation of copper cells (Wang et al., 2014), a specialized cell type that attenuates bacterial growth by establishing a stomach-like region of low pH (Li et al., 2016). Thus, we speculate that *Notch*-deficient intestines are impaired in their ability to eradicate bacteria due to a multifactorial network of perturbed immunity and mis-differentiation of mature epithelial cells.

To determine how *Lb* affects intestinal growth, we characterized transcriptional responses from midgut progenitors of flies inoculated with *Lb*. We observed significant effects of *Lb* on expression and subcellular localization of integrins, critical regulators of stem cell-niche interactions (Ellis and Tanentzapf, 2010; Fernandez-Minan et al., 2007; Marthiens et al., 2010; Toyoshima and Nishida, 2007), and stem cell maintenance (Goulas et al., 2012; Lin et al., 2013; Okumura et al., 2014; You et al., 2014). Typically, integrins accumulate at basolateral margins of intestinal progenitors, and anchor interphase progenitors to the extracellular matrix. In many tissues, including the fly intestine, integrins organize the stem cell division plane, by orienting the mitotic spindle. Progenitors mainly divide at angles greater than 20° to the basement membrane, leading to asymmetric divisions (Hu and Jasper, 2019; Ohlstein and Spradling, 2007), where basal daughter cells remain in the niche and retain stemness, while apical daughters exit the niche and differentiate. In contrast to other tissues, approximately 20% of divisions in the young adult intestine occur symmetrically, yielding clonal lineages of stem cells or enteroblasts (Hu and Jasper, 2019; Jin et al., 2017; O’Brien et al., 2011). Over time, enteroblast clones differentiate into mature enterocytes that eventually die. However, clonal stem cell lineages retain the capacity to grow and establish regional dominance within the epithelium. In some cases, symmetric divisions facilitate adaptive responses to environmental fluctuations allowing the intestinal environment to tune proliferative needs to extrinsic factors.

For instance, rapid changes in nutrient availability, or ingestion of toxic doses of paraquat increase the frequency of symmetric divisions and expand the stem cell pool, allowing for a rapid regenerative response (Hu and Jasper, 2019; O’Brien et al., 2011). We discovered that *Lb* disrupts subcellular distribution of integrins in progenitors, and promotes symmetric expansion of ISCs. *Lb*-mediated ISC growth likely accounts for the tumor burden noted in Notch-deficient flies inoculated with *Lb*. However, as individual stem cell lineages accumulate mutations with age, we consider it possible that exposure to *Lb* also facilitates spontaneous development of intestinal tumors in wild-type adult flies. Since our focus was on intestinal progenitors it is unclear whether *Lb* also alters integrins in mature epithelial cells. As integrin loss in enterocytes causes delamination and subsequent stress induced tumor growth (Patel et al., 2015), it would be pertinent to understand whether the actions of *Lb* on integrins is specific to progenitors or if enterocytes are also affected.

How *Lb* disrupts integrins and promotes stem cell growth requires clarification. As stem cells derive cues from the surrounding epithelium to direct their growth, we consider it likely that mature epithelial cells, such as enterocytes, sense *L. brevis* and transduce signals to ISCs to promote growth. Along these lines, *Lp* and *Lb* induce ROS production in enterocytes leading to increased intestinal mitoses (Jones et al., 2013; Jones et al., 2015; Lee et al., 2013; Reedy et al., 2019). Interestingly, the cell wall of *Lb* is sufficient for tumor growth, suggesting a role for bacterial recognition pathways in intestinal growth. Consistent with this hypothesis, constitutive activation of the IMD pathway promotes growth of Notch-deficient tumors (Petkau et al., 2017), and chronic inflammation is a risk factor for development of colorectal cancer (Kim and Chang, 2014). Therefore, we speculate that *Lb* activates immune programs such as ROS and IMD in the epithelium, which decreases integrin expression in ISCs, promoting symmetric expansion of stem cells and increasing the regenerative capacity of the wildtype intestine, but also driving growth of mutant progenitors. Due to the evolutionary conservation of intestinal homeostatic regulators, we believe *Drosophila* will be a fruitful model to determine how gut-resident bacteria influence intestinal progenitor function.

## ACKNOWLEDGEMENTS

We thank Dr. Bruce Edgar, Dr. Bruno Lemaitre and Dr. Lucy O’Brien for providing fly lines. We acknowledge microscopy support from Dr. Steven Ogg, Gregory Plummer and Woo Jung Cho at the Faculty of Medicine and Dentistry Imaging core; flow cytometry support from Dr. Aja Rieger at the Faculty of Medicine and Dentistry Flow Cytometry Core; and cryo-sectioning support from Lynette Elder at the Alberta Diabetes Institute Histocore. The authors wish to thank Kin Chan at the Network Biology Collaborative Centre (nbcc.lunenfeld.ca) for the RNA-Seq service. Network Biology Collaborative Centre is a facility supported by Canada Foundation for Innovation, the Ontarian Government, and Genome Canada and Ontario Genomics (OGI-139). This work was supported by grants from the Canadian Institute of Health Research (Grant # PJT 159604). Anthony Galenza and David Fast have funding support through the National Science and Engineering Research Council scholarships. Meghan Ferguson has funding through Alberta Innovates Graduate Student Scholarships.

## MATERIALS AND METHODS

### *Drosophila* husbandry

*Drosophila* stocks and crosses were setup and maintained at 18-25°C on standard corn meal food (Nutri-Fly Bloomington formulation; Genesse Scientific) with a 12:12 light:dark cycle. All experimental flies were virgin female flies except when noted. Upon eclosion, flies were kept at 18°C then shifted to the appropriate temperature once 25-30 flies per vial was obtained. Germ free and mono-associated flies were generated as previously described (Fast et al., 2018). To generate germ free flies, freshly eclosed flies were fed autoclaved food with antibiotics (Ampicillin (100μg/mL), Neomycin (100μg/mL), Vancomycin (50μg/mL) and Metronidazole (100μg/mL)) for 5 days at 25°C. Conventionally reared controls were fed autoclaved food without antibiotics for 5 days at 25°C. To generate monoassociated animals, flies were made germ free as above then were fed 1mL OD_600_=50 of *L. brevis* or *L. plantarum* resuspended in sterile 5% sucrose/PBS on a cotton plug overnight at 25°C. During this overnight feeding, CR and GF controls were fed sterile 5% sucrose/PBS without bacteria. The following morning, CR, *Lb* and *Lp* conditions were transferred to fresh autoclaved food at 29°C for the remainder of the experiment, while GF flies were maintained on autoclaved food with antibiotics. Sterility and mono-association were confirmed in each experimental vial by plating fly homogenate or fly food on MRS. Vials found to be contaminated were discarded from the experiment. Fly lines used in this study were: *w;esg-GAL4,tubGAL80^ts^,UAS-GFP* (*esg^ts^)*, *UAS-N^RNAi^* (VDRC ID# 100002), w;*Myo1A-GAL4;tubGAL80^ts^,UAS-GFP (Myo1A^ts^)*, *imd*, *w^1118^*, *UAS-ImdCA* (Petkau et al., 2017), *w;esg-GAL4,UAS-CFP, Su(H)-GFP;tubGal80^ts^ (esg^ts^,UAS-CFP,Su(H)-GFP)*. Twin-spot MARCM flies were generated by crossing together *hsFLP;FRT40A,UAS-CD2-RFP,UAS-GFP-miRNA/cyo* flies and *FRT40A,UAS-CD8-GFP,UAS-CD2-miRNA/cyo;tubGAL4/TM6C* flies to create *hsFLP;FRT40A,UAS-CD2-RFP,UAS-GFP-miRNA/FRT40A,UAS-CD8-GFP,UAS-CD2-miRNA;tubGAL4/+.* Twin-spot clones were induced by a 37°C heatshock for 1 hour immediately after transferring flies to fresh food after the overnight monoassociation protocol. Flies were then shifted back to 29°C for 8 days before dissecting and counting clones from 30 intestines per treatment. The *imd* and *UAS-ImdCA* lines had been backcrossed into our wild-type *w^1118^* background.

### Bacterial strains and growth conditions

*L. brevis^EF^* (DDBJ/EMBL/GeneBank accession LPXV00000000) and *L. plantarum^KP^ (*DDBJ/EMBL/GenBank chromosome 1 CP013749 and plasmids 1-3 CP013750, CP013751, and CP013752) were both isolated from our *Drosophila* lab stocks and have been previously described (Petkau et al., 2016). Both bacteria were streaked out on MRS plates (BD; 288210) and aerobically grown at 29°C. Single colonies were picked for growth in MRS broth (Sigma; 69966) at 29°C (*L. brevis* for 2 days and *L. plantarum* for 1 day). To generate dead *L. brevis*, liquid culture was spun down, washed twice with sterile water then resuspended in sterile water before heating to 95°C for 30min. After heating, the killed bacteria was spun again and resuspended to 10mg/mL in sterile 5% yeast, 5% sucrose in PBS.

To extract the cell wall, *L. brevis* was heat killed as above, let cool on ice, then run through a French Press at 20,000 psi three times to lyse the bacterial cells. After lysis, any remaining whole cells were collected and discarded by two successive spins at 2000g. To collect the cell wall, the supernatant was spun at 10,000g for 30 minutes, washed twice with 1M NaCl and twice with sterile water before resuspending in sterile 5% yeast, 5% sucrose in PBS. Germ free flies were fed a 10mg/mL cell wall solution on filter paper disks on top of sterile food alongside 10mg/mL dead *L. brevis* and sterile 5% yeast, 5% sucrose PBS without any *L. brevis* extracts. Dead bacteria and cell wall were continuously fed to flies during the experiment, with fresh extracts provided every second day. Sterility of dead *Lb* and cell wall was confirmed by plating 100uL of extract on MRS.

### Immunofluorescence

Intestines were dissected in PBS, fixed in 4% formaldehyde for 20 minutes then blocked overnight at 4°C in 5% normal goat serum (NGS), 1% bovine serum albumin (BSA), and 0.1% tween-20. Washes were done in blocking solution without NGS. Primary and secondary antibody incubations were done for 1 hour at room temperature in blocking buffer without NGS. For Delta and PH3 stain, we used a revised protocol where 8% formaldehyde was used to fix, washes were done in PBS with 0.2% TritonX-100 (PBST) and blocked in PBST with 3% BSA. To prepare the intestinal sections, the posterior midgut was extracted, flash frozen on dry ice in frozen section compound (VWR 95057-838), sectioned to 10μm and slides were stained using the same parameters as whole guts. Primary antibodies used: anti-prospero (1/100; Developmental Studies Hybridoma Bank(DSHB)), anti-mys (1/100; DSHB), anti-GFP (1/1000; Invitrogen), anti-phospho-histone3 (1/1000; Upstate), anti-Delta (1/100; DSHB). Secondary antibodies used: goat anti-mouse 568 (1/500; Invitrogen), goat anti-rabbit 488 (1/500; Invitrogen). DNA stains used: Hoechst (1/500; Molecular Probes), DRAQ5 (1/500; Invitrogen). Apoptotic cells were detected in dissected guts using the TMR red In Situ Cell Death Detection Kit (Roche; 12156792910) following standard kit staining protocol. GFP primary antibody was used for the TUNEL experiment and cryosectioning experiments to retain GFP signal from the esg promoter. Intestines were mounted on slides using Fluoromount (Sigma; F4680). For every experiment, images were obtained of the posterior midgut region (R4/5) of the intestine with a spinning disk confocal microscope (Quorum WaveFX). PH3+ cells were counted through the entire midgut. To determine the apical and basal mys intensity, we drew a line of 10 pixel width from the basal side to the apical (lumen side) side across GFP+ progenitor cells. We defined apical and basal progenitor cell borders as 50% of the maximum GFP intensity, as this GFP intensity coincides with the basal mys intensity peak. We determined the intensity of GFP and mys across the progenitors using the function plot profiles, copied these values into Excel and determined the apical and basal mys intensities. All image stacking, intensity and area calculations were done using Fiji software (Schindelin et al., 2019).

### Intestinal progenitor cell isolation and RNA sequencing

Progenitor cell isolation by fluorescence activated cell sorting (FACS) was adapted from previously described protocols (Dutta et al., 2013). In brief, three biological replicates consisting of 100 fly guts per replicate with the malphigian tubules and crop removed were dissected into DEPC PBS and placed on ice. Guts were dissociated with 1mg/ml of elastase at 27°C with gentle shaking and periodic pipetting for 1hour. Progenitors were sorted based on GFP fluorescence and size with the BD FACSAria III sorter. All small GFP positive cells were collected into DEPC PBS. Cells were pelleted at 1200g for 5 minutes and then resuspended in 500μl of Trizol. Samples were stored at −80°C until all samples were collected. RNA was isolated via a standard Trizol chloroform extraction and the RNA was sent on dry ice to the Lunenfeld-Tanenbaum Research Institute (Toronto, Canada) for library construction and sequencing. The sample quality was evaluated using Agilent Bioanalyzer 2100. TaKaRa SMART-Seq v4 Ultra Low Input RNA Kit for Sequencing was used to prepare full length cDNA. The quality and quantity of the purified cDNA was measure with Bioanalyzer and Qubit 2.0. Libraries were sequenced on the Illumina HiSeq3000 platform.

### RNA Sequencing data processing and analysis

On average, we obtained 30 million reads per biological replicate. FASTQC was used to evaluate the quality of raw paired-end sequencing reads (http://www.bioinformatics.bbsrc.ac.uk/projects/fastqc, version 0.11.3). Adaptors and reads of less than 36 base pairs in length were trimmed from the raw reads using Trimmomatic (version 0.36) (Bolger et al., 2014). Reads were aligned to the *Drosophila* transcriptome- bdgp6 (https://ccb.jhu.edu/software/hisat2/index.shtml) with HISAT2 (version 2.1.0) (Kim et al., 2015). The resulting BAM files were converted to SAM flies using Samtools (version 1.8) (Li et al., 2009). The converted files were counted using Rsubread (version 1.24.2) (Liao and Smyth, 2019) and loaded into EdgeR (version 3.16.5) (Robinson et al., 2010). In EdgeR, genes with counts less than 1 count per million were filtered and libraries were normalized for size. Normalized libraries were used to call genes that were differentially expressed among treatments. Genes with P-value < 0.01 and FDR < 5% were defined as differentially expressed genes.

Principle component analysis was performed on normalized libraries using Factoextra (version 1.0.5). Gene Ontology enRIchment anaLysis and visuaLizAtion tool (GORILLA) was used to examine Gene Ontology (GO) term enrichment(Eden et al., 2009). Specifically, differentially expressed genes (defined above) were compared in a two-list unraked comparison to all genes output from edgeR as a background set. Redundant GO terms were removed.

### Quantification of Bacterial CFUs

Five flies were randomly selected from a single vial of flies for each biological replicate and surface sterilized by washing in 10% bleach and 70% ethanol. Flies were then homogenized in MRS, serially diluted and 10μL of each dilution was plated on MRS. Colonies were counted from 10μL streaks that had 10-200 colonies and the CFU/fly calculated.

### TEM

Intestines were dissected from virgin female flies that had been at 29°C for 8 days following germ free and bacterial association protocols. Posterior midguts were excised and fixed with 3% paraformaldehyde with 3% glutaraldehyde. Fixation, contrasting, sectioning, and visualization were performed at the Faculty of Medicine and Dentistry Imaging Core at the University of Alberta. Midgut sections were visualized with Hitachi H-7650 transmission electron microscope at 60Kv in high contrast mode.

### Lifespan

Virgin female flies were monoassociated with *L. brevis* or raised with a conventional microbiome as described. After mono-association, flies distributed to sterile vials with autoclaved food at a density of 10 flies/vial shifted to 29°C for the remainder of the experiment. Dead flies were counted every 1-3 days and vials were flipped 3 times per week to fresh autoclaved food.

### Data visualization and Statistical analysis

Figures were constructed using R (version 3.3.1) via R studio (version 1.1.442) with easyggplot2 (version 1.0.0.9000), with the exception of GO term figures and lineplots where ggplot2 (version 3.0.0) was used. Longevity graphs were made in Prism software along with the stats. All other statistical analysis was performed in R. Figures were assembled in Adobe Illustrator.

### Data availability

Gene expression data has been deposited to the NCBI GEO database accession GSE138555.

## SUPPLEMENTARY FIGURES

**Supplementary Figure 1.**
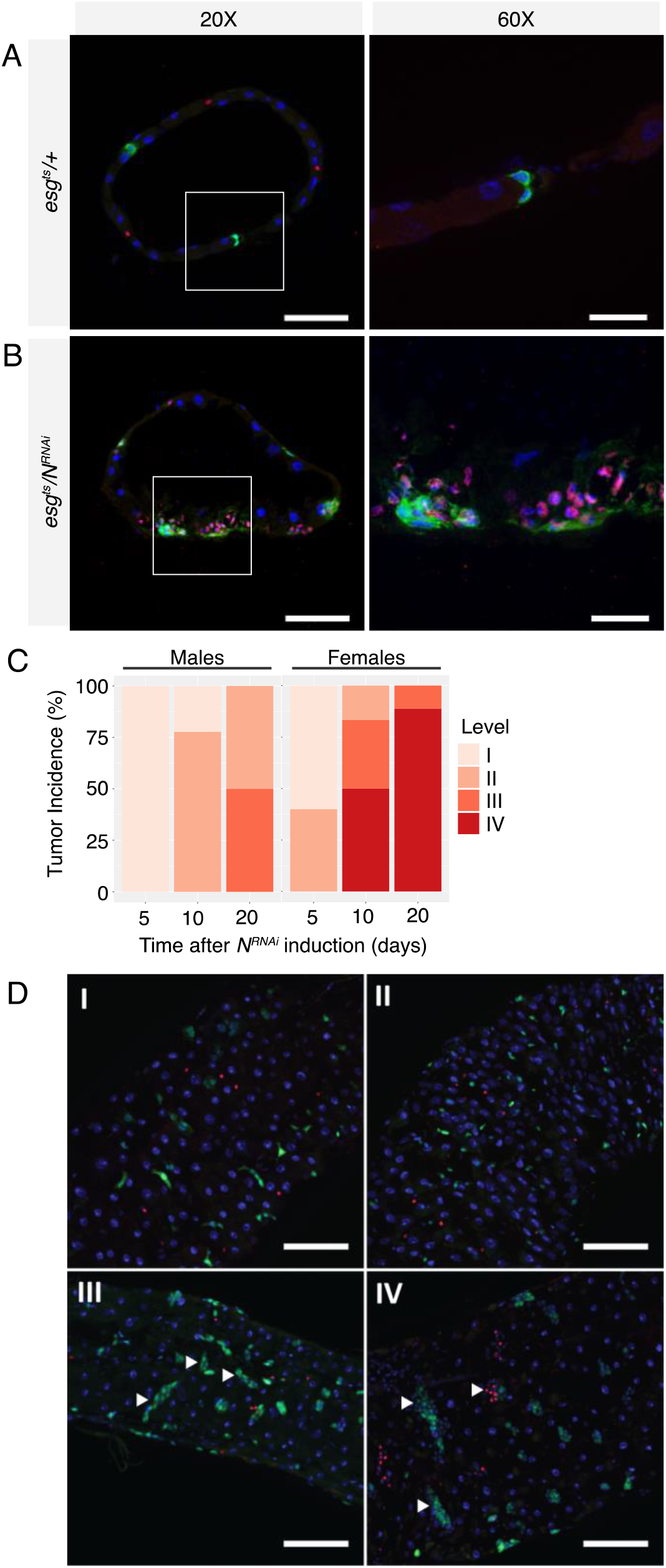
Multilayered *Notch*-deficient tumors form in female *Drosophila* intestines. **(A)** Cross section of *esg^ts^/+* posterior midgut and **(B)** cross section of *esg^ts^/N^RNAi^* showing multilayered tumors composed of enteroendocrine cells labelled by Prospero (red) and progenitors (green). DNA labelled with Hoechst (blue). Scale bars: 20X = 50μm 60X = 15μm. **(C)** Incidence of tumors in male and virgin female *esg^ts^/N^RNAi^* intestines 5, 10 and 20 days after *Notch* knockdown in intestinal progenitors. **(D)** Tumor incidence grading system. I – healthy intestine, II – intestinal dysplasia without tumorigenesis, III – tumors populated by progenitors, IV – tumors populated by progenitors and enteroendocrine cells.

**Supplementary Figure 2.**
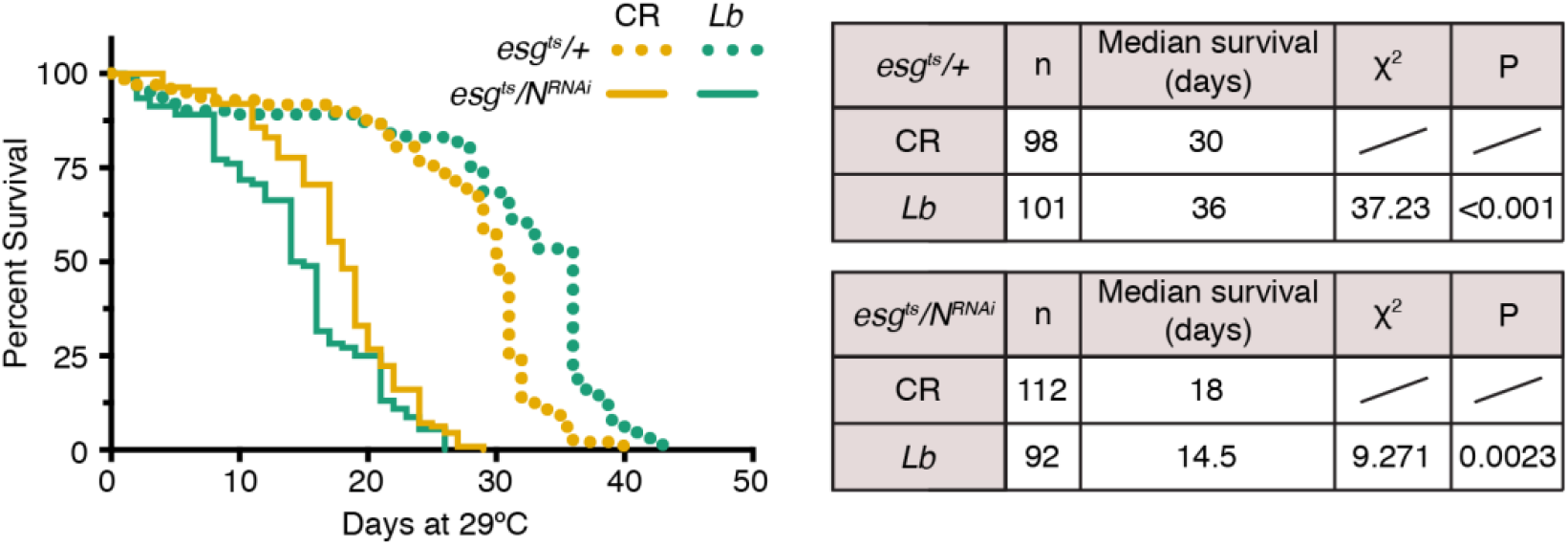
*L. brevis* reduces the lifespan of flies upon intestinal Notch-inactivation. Fly lifespan of CR (yellow) and Lb colonized (teal) *esg^ts^/+* and *esg^ts^/N^RNAi^* flies was monitored after switching to the permissive temperature of 29°C. Statistical analysis performed in Prism software using Log-rank Mantel-Cox test.

**Supplementary Figure 3.**
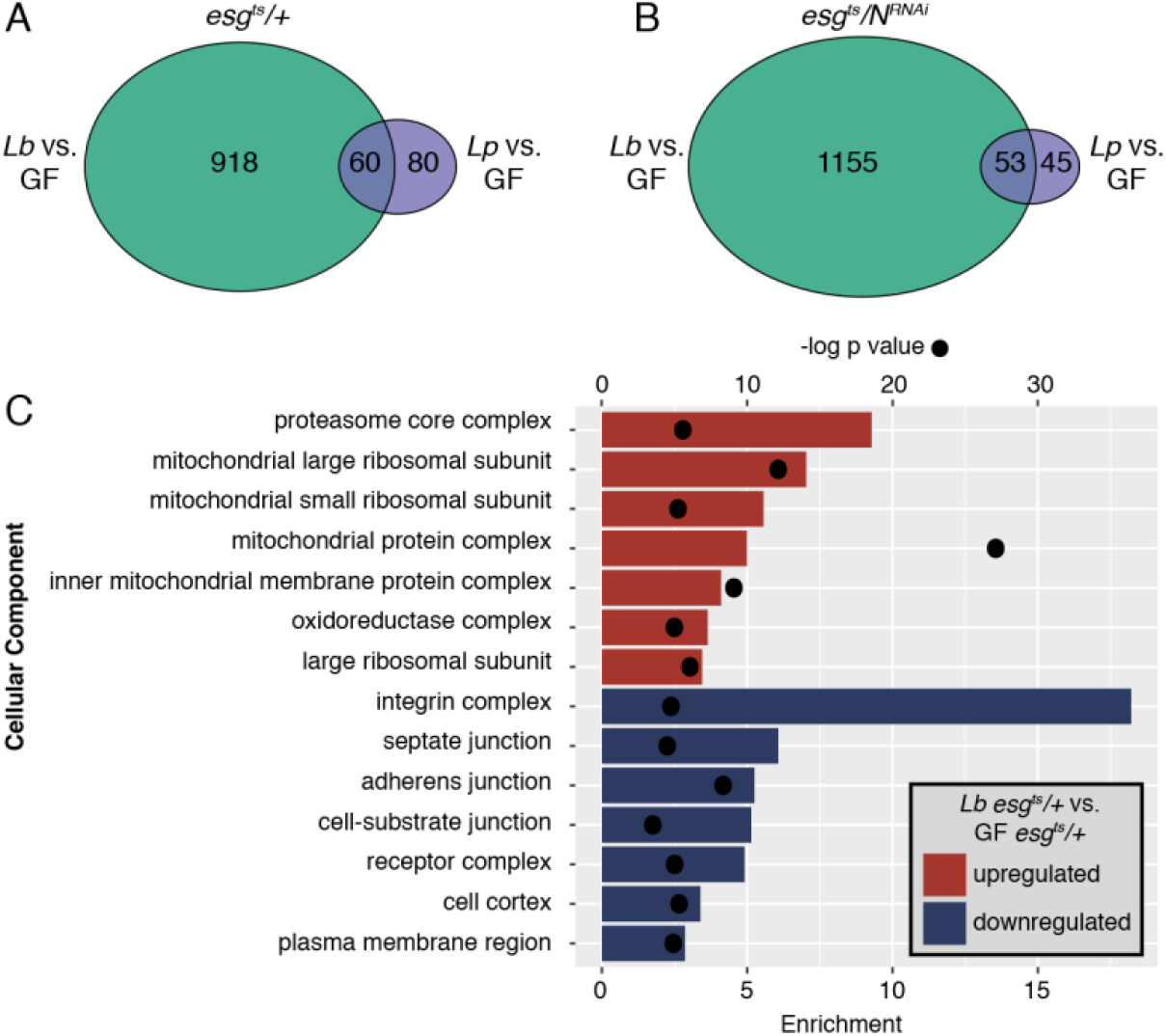
Specific effect of *L. brevis* on integrin expression. **(A-B)** Number of significant genes (p<0.01, FDR<5%) differentially expressed in progenitors upon *Lb* or *Lp* mono-association in comparison with GF. **(A)** Comparisons from *esg^ts^/+*. **(B)** Comparisons from *esg^ts^/N^RNAi^*. **(C)** Cellular component GO terms enriched in progenitors from *esg^ts^/+* colonized with *Lb* compared to GF. Enrichment score shown as bars and p values shown as dots.

**Supplementary Figure 4.**
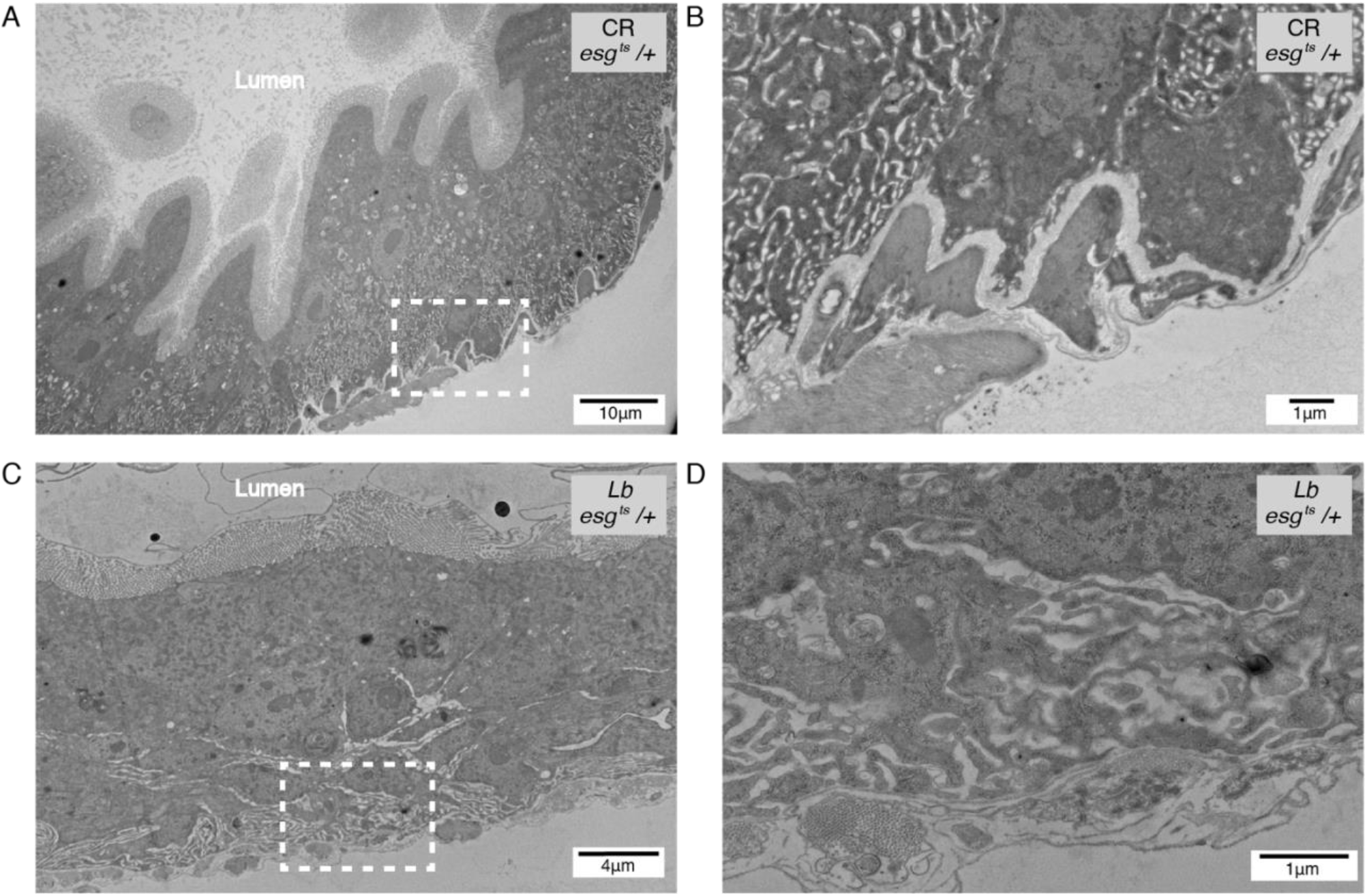
Colonization with *L. brevis* disrupts basal morphology in the posterior midgut. **(A,B)** TEM images of a CR *esg^ts^/+* posterior midgut highlighting easily discernable extracellular matrix and visceral muscle in close contact with an intestinal progenitor cell. **(C,D)** TEM images of a *Lb* mono-associated *esg^ts^/+* posterior midgut highlighting basal disruption. Dashed boxes in (A) and (C) showed zoomed areas of (B) and (D) respectively.

**Supplementary Figure 5.**
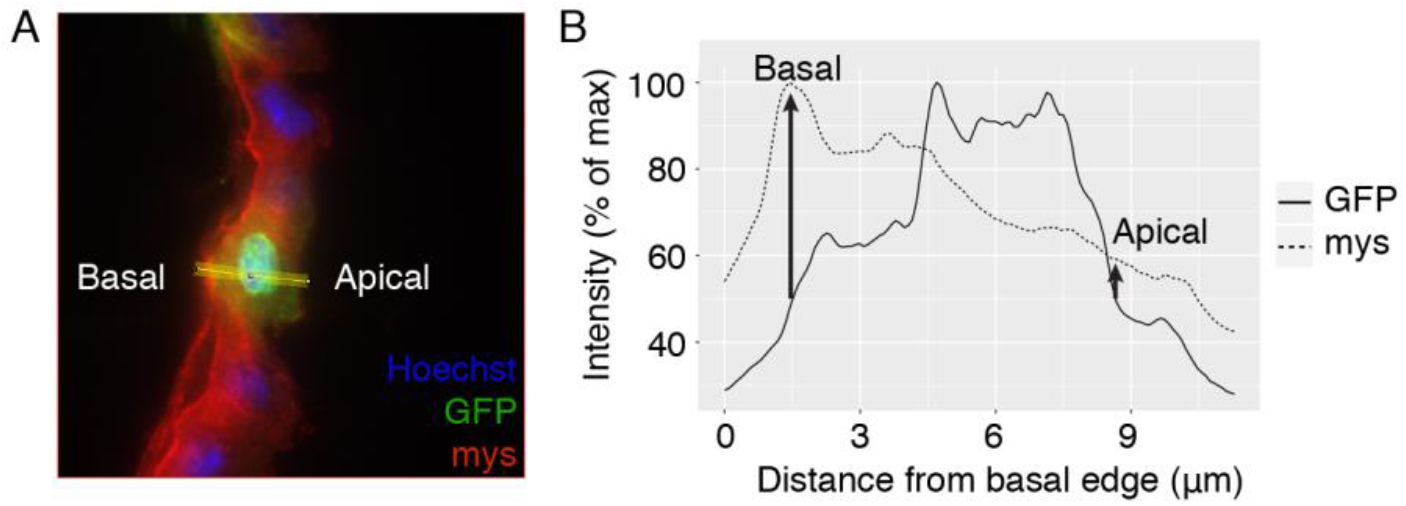
Analysis of apicobasal integrin localization. **(A)** Example image of intestinal cross section where intensities of mys (red) and GFP (progenitors) were measured along the yellow line from basal to apical edges of progenitors. **(B)** Plot showing the normalized intensities of GFP (solid line) and mys (dashed line) across the yellow line in (A). The basal and apical edges of progenitors were defined as 50% of the maximum GFP intensity and mys intensity was determined at each of these edges.

**Supplementary Figure 6.**
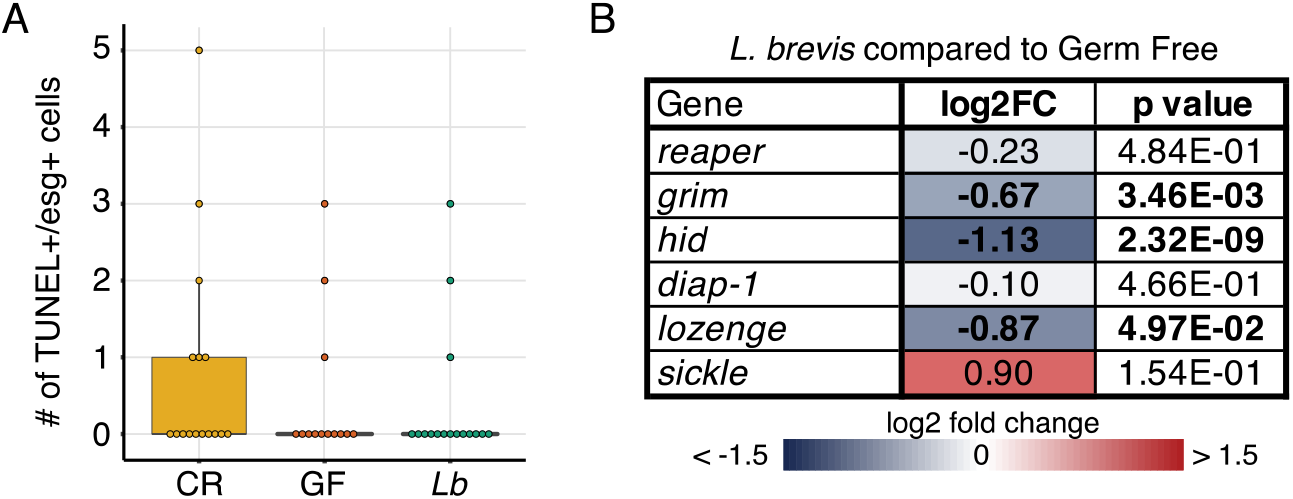
*L. brevis* does not enhance progenitor cell death. **(A)** Total number of TUNEL positive progenitor cells in the posterior midgut of CR, GF and *Lb* monoassociated *esg^ts^/+* flies. **(B)** Apoptotic gene expression from FACS isolated intestinal progenitor cells in response to *Lb* colonization. Bolded values are significant at p<0.05, FDR <5%.

## Notes

### Competing Interest Statement

The authors have declared no competing interest.

### Summary of Updates

This revised version includes additional data where we show that Lactobacillus brevis increases the rate of symmetric intestinal stem cell divisions in the adult fly gut.

